# Mixture of Experts for Predicting Antibody-Antigen Binding Affinity from Antigen Sequence

**DOI:** 10.1101/511360

**Authors:** Srivamshi Pittala, Chris Bailey-Kellogg

## Abstract

Antibodies provide a key mode of defense employed by the immune system to fight disease, so eliciting potent antibodies is one of the main goals in vaccine development. Antibodies are also being conceived as powerful therapeutic agents, so engineering potent antibodies is one of the main goals in biologic drug development. The power of antibodies lies in their affinity and specificity in recognizing their cognate antigens. Unfortunately, experimental techniques to determine antibody-antigen binding affinities are difficult to scale up to large sets of new antibodies and antigens. Though computational methods are suitable for large-scale prediction, current methods lack sufficient accuracy. Here, we address the problem of predicting the binding affinity of an antibody against an antigen variant based on the amino acid sequence of that antigen variant. We develop a mixture of experts approach that learns models for individual antibodies against some antigen variants, and then combines information across the antibodies in order to make accurate predictions for a wide range of new variants. In evaluation on a dataset consisting of 52 antibodies and 608 strains of HIV, the predictive accuracy of our approach is demonstrated to be significantly better than that of an existing approach. Our method provides a fast and accurate way to predict antibody-antigen binding affinity, which has the potential to expedite the study of antibody-antigen interactions for vaccine design and therapy.

## 1 Introduction

Recognition and binding of antigens by antibodies is one of the primary processes by which our immune system protects us from infections [1]. Therefore, studying antibody-antigen interactions to understand the factors driving the specificity and affinity of antibodies to their antigens has the potential to advance vaccine development [2]. Furthermore, recombinant antibodies are being employed as therapeutics, and knowledge of antibody mechanism of action can help guide better design and development [3]. Binding affinity measurements between antibodies and antigens thus offer means to analyze antibody-antigen interactions [4]. Experimental techniques to determine binding affinities are resource intensive and time consuming to scale-up to newly discovered antigens and antibodies, therefore necessitating the development of computational methods to predict antibody-antigen binding affinity [5].

In this work, we focus on predicting binding affinities of antibodies against antigens that exhibit great variation to escape from the immune system, like HIV. Development of an effective vaccine/treatment against HIV is challenging in part due to the virus’; high rate of mutation allowing to escape from immune recognition [6]. Therefore one of the main goals in HIV vaccine and therapeutic antibody development is the elicitation, discovery, and engineering of broadly neutralizing antibodies (bnAbs) that neutralize widely differing variants of the virus. Some bnAbs have been discovered in HIV infected individuals [7] and are now being pushed toward the clinic [8]. Binding affinity measurements between bnAbs and HIV variants have been used to understand the evolution of bnAbs to neutralize multiple strains of HIV [9], and also to identify binding sites on the virus [10, 11, 12]. However, it is infeasible to experimentally determine binding affinities between bnAbs and variants of the rapidly mutating virus. Recently, a learning-based computational approach was proposed for antibody-specific prediction of binding affinity to new viral strains [11]. Their approach focused on identifying antibody-specific binding sites on the surface of HIV, and was constrained to using only binding affinity data corresponding to the antibody of interest. However, we surmise that binding affinity patterns across multiple antibodies can be leveraged for the same task of antibody-specific binding affinity prediction. To this end, we propose a mixture of experts based approach to predict binding affinity of antibodies to new strains, taking as input only the protein sequence of the strains. Our results show a significant boost in accuracy with our approach compared to the existing approach.

## 2 Methods

Given a sparse binding affinity matrix 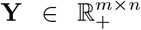 between a set of *m* antibodies **A** = {*a*_1_, *a*_2_,…, *a*_m_} and a set of *n* antigens **V** = {*v*_1_, *v*_2_,…, *v*_*n*_}, the objective is to perform antibody-specific (row-wise) completion of the matrix. Matrix completion is a well known problem with applications to recommender systems [13, 14], where the goal, for example, is to predict a user’;s interests based on similarity to other users’; interests. We apply a similar strategy here to predict binding affinities of an antibody by leveraging binding affinity patterns from multiple antibodies. Particularly, we use a mixture of experts model [15], where a mixture model to predict binding affinity of an antibody is learned to combine the predictions from multiple antibodies (experts). We employ a two step approach to learn a mixture of experts model for each antibody. First, an expert model **E**_*i*_ corresponding to antibody *a_i_* is learned based its binding affinity values 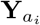. Next, a mixture model **M**_*i*_ for an antibody *a*_*i*_ is learned by combining the expert models of the other antibodies *a*_*j*:*j*≠*i*_.

### 2.1 Learning Antibody Experts

An individual antibody expert model is learned following the formulation of the problem of predicting binding affinity *y*_(*a*,*v*)_ between a given antibody *a* and an antigen *v* as a *l*_1_-regularized linear regression task [11]. In this formulation, the antigens are input as aligned amino acid sequences represented by applying one-hot encoding, where each residue of the sequence is encoded as a binary vector indicating one of the 21 (‘gap’ included) amino acids present at that position. For the antibody *a*, the expert model *E* is learned by minimizing the regularized squared error between the observed binding affinity values **Y**_*a*_ and the model’s prediction (Eqn 1).

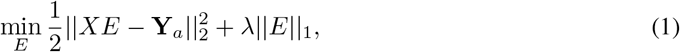

where *X*_*k*×*p*_ denotes the feature matrix consisting of *k* antigens, each of length *p* after alignment and binary representation. 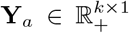 is the vector of observed binding affinity values. The regularization parameter *λ* is selected by performing a 10-fold cross-validation, and choosing the value with lowest mean squared error averaged across folds.

### 2.2 Learning Mixture of Antibody Experts

The expert models are learned using only binding affinity data corresponding to an antibody, and hence represent antibody-specific signatures of interaction with the antigens. We propose to learn a mixture model to leverage interaction signatures from multiple antibodies. We formulate the predicted binding affinity *y*_(*a_i_*,*v*)_ between a given antibody *a*_*i*_ and a viral strain *v*, as a linear combination of the predictions of other expert models *E*_*j*:*j≠i*_ for *v* [15]. The gating function (i.e., weights of the combination) is learned specific to the antibody *a_i_*. For the antibody *a_i_*, a mixture model *M_i_* is learned by minimizing the squared error between the observed binding affinity values 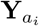 and the model’;s prediction (Eqn 2). We apply *l*_1_-regularization here to use only a few of the relevant antibody models for prediction.

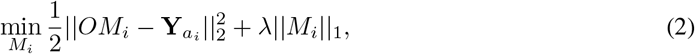

where *O* ∈ ℝ^*k*×(*m*−1)^ denotes the feature matrix, each column of which is the prediction of the other *m* − 1 expert models on 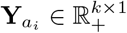 is the vector of observed binding affinity values. The regularization parameter *λ* is again selected by performing 10-fold cross-validation.

## 3 Experimental Setup

To evaluate the prediction performance of our approach, we used the publicly available CATNAP database [16], which has neutralization measurements for about 200 antibodies against a panel of about 1000 HIV strains (about 18% complete). In this dataset, neutralization measurements were reported as minimum inhibitory concentration of the antibody needed to reduce the infectivity of the virus by 50% (IC_50_). IC_50_ measurements above 50 *μg/ml*. were considered to indicate no neutralization
of the antibody by the database providers, and hence those measurements were truncated to 50 *μg/ml*. As in the previous studies using neutralization titers, IC_50_ values were log_10_ transformed to be considered as binding affinity measurements. Since our approach focuses on antibody-specific binding affinity prediction, we eliminated antibodies with fewer than 45 neutralized antigens, to ensure there would be sufficient testing cases for validation. This step resulted in a smaller dataset of 52 antibodies and 608 strains (about 35% complete). The feature matrices were column normalized to have zero mean and unit variance before learning the expert and mixture models for each antibody. The prediction performance of the models was tested via 10-fold cross-validation, using Pearson correlation coefficient (PCC) between the observed and predicted affinity values as the metric.

## 4 Results

We compare the prediction performance of the individual expert models [11] and our proposed mixture models in Figure 1. The mixture models, with mean PCC of 0.68 across the 52 antibodies, substantially outperform the individual models, which have a mean PCC of 0.48. Figure 1B shows that the increase in performance when using the mixture models is also statistically significant. Thus, this result demonstrates that the task of predicting antibody-antigen binding affinity from antigen sequences clearly benefits from combining multiple antibody-specific signatures of interaction with the antigens.

**Figure 1:**
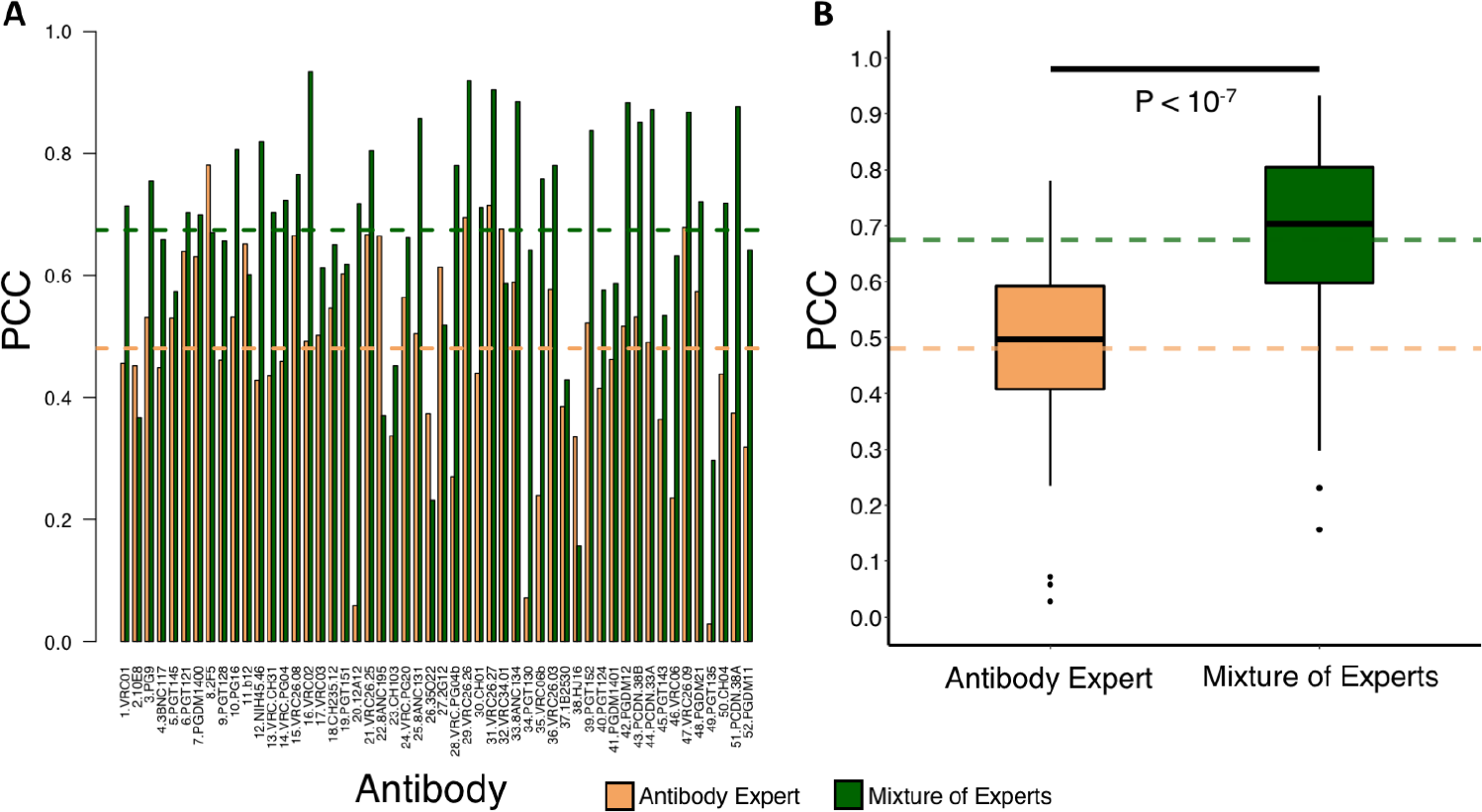
(A) Per-antibody PCC between the observed and predicted binding affinities of the expert (yellow) and mixture (green) models learned using all antigens. (B) Box-plot comparison of performance across 52 antibodies, with Wilcoxon-Mann-Whitney test. Horizontal dashed lines represent mean PCC across 52 antibodies.

To check whether mixture models were dominated by a few antibodies, we counted the number of times each antibody’;s expert model contributed to one of the 52 mixture models. As shown in Figure 2A, the distribution is fairly uniform, with no clear signs of the mixture models being dominated by a few expert models. In addition, we see in Figure 2B that frequency of appearance of expert models in the mixture models shows no strong correlation with the similarity in antibody sequence. The top three most frequent antibodies are positioned far from each other.

Furthermore, we sought to evaluate the utility of sequence similarity of antibodies in training mixture models by replacing the learned weights of the gating function with antibody sequence similarity distances. Particularly, we used 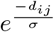 as the weight to combine the prediction output of the expert model *E*_*j*:*j*≠*i*_, where *d*_*ij*_ is the distance between heavy chains of antibody *a*_*i*_ and *a_j_* [17], and *σ* is the tuning parameter. The predictions made by such sequence based mixture models showed poor correlation to observed binding affinities. In fact, their mean PCC across the 52 antibodies was 0.35, which is significantly lower than that of both expert models’ 0.48 and mixture models’ 0.68 reported above.

**Figure 2:**
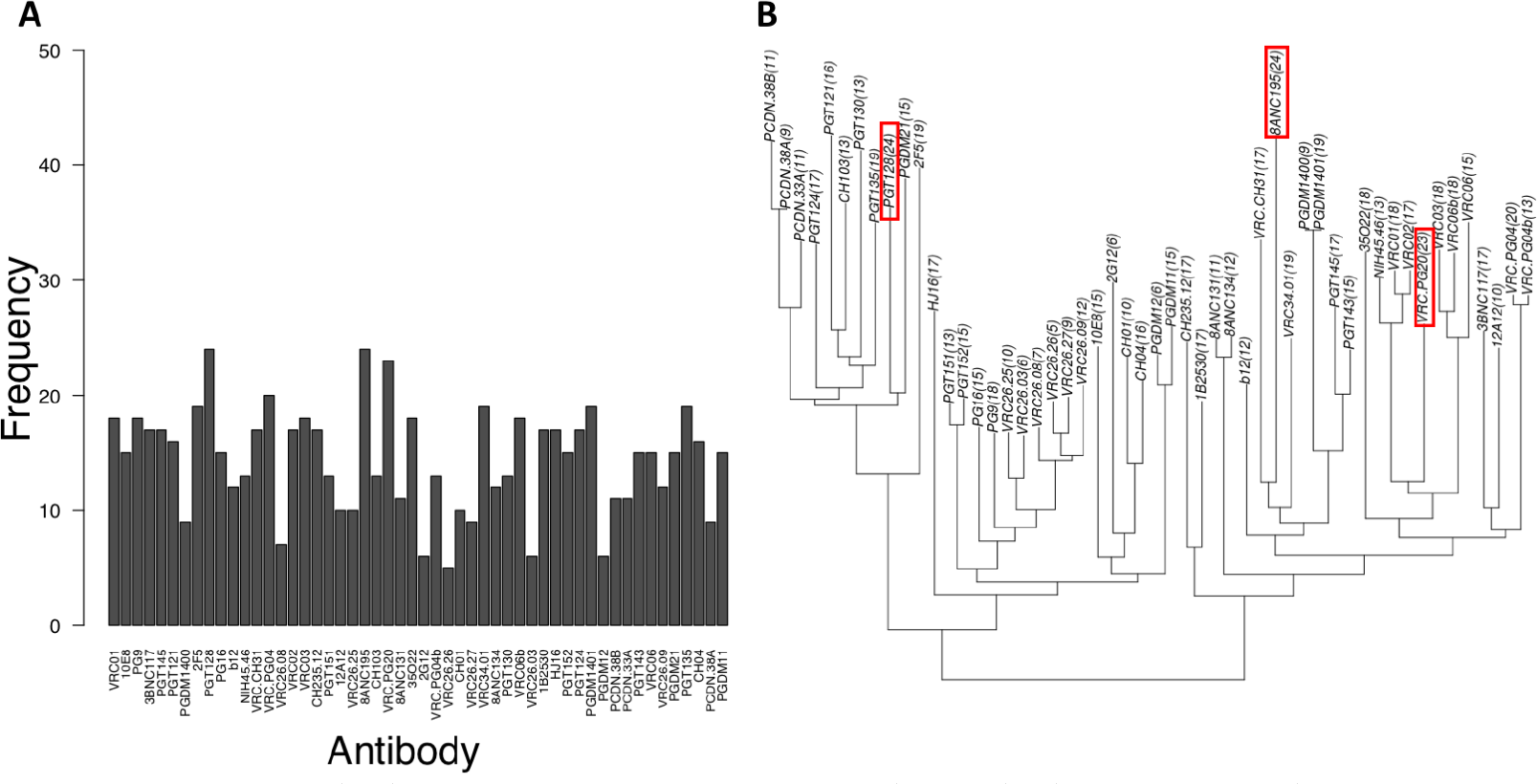
(A) Per-antibody frequency of appearance of the antibody’s expert model in 52 mixture models. (B) Dendrogram of antibodies clustered based on their heavy chain protein sequences with their frequencies shown in parentheses. The top three most frequent antibodies are highlighted in red.

Since the IC_50_ values of non-neutralized antibody-antigen pairs were truncated to 50 *μg/ml*, it is possible that the learning objective is overwhelmed by constant-valued samples. Hence, we evaluated the performance of the models learned using only neutralized antigens (IC_50_< 50 *μg/ml*). The prediction performance of both expert and mixture models dropped, with the mean PCC of 0.23 and 0.50 respectively, compared to when using all the antigens (Figure 1A). This result shows that both non-neutralized and neutralized antigens are beneficial to predicting antibody-antigen interaction. This result again illustrates the benefit of using mixture models, which significantly outperform the expert models.

## 5 Conclusions

We have presented a mixture of experts based learning approach to predict antibody-antigen binding affinity solely from the antigen sequence. Building on an existing approach that learns antibody-specific expert models, we learn a gating function to effectively combine binding affinity predictions of expert models corresponding to different antibodies. Our results show that the mixture models achieve significantly higher accuracy compared to the individual expert models, providing a better way to predict binding affinity between existing antibodies and new antigens. Though we applied our approach to the case of HIV, it can be applied to other infectious diseases, particularly where a diverse set of antigens are involved. This work could be further extended to use a combined classification and regression approach that simultaneously predicts a non-binding versus binding interaction as well as the binding affinities of binding antibodies. Finally, it will be interesting to study if such models could also be used to identify structural sites of antibody-antigen interaction.

## 6 Acknowledgements

We thank Dr. Qiang Liu for help with initial formulations of the prediction problem. This work was supported in part by NIH grant R01-GM-098977. We also gratefully acknowledge computational resources provided by NSF grant CNS-1205521.

